# Learning probabilistic protein-DNA recognition codes from DNA-binding specificities using structural mappings

**DOI:** 10.1101/2022.01.31.477772

**Authors:** Joshua L. Wetzel, Kaiqian Zhang, Mona Singh

## Abstract

Knowledge of how proteins interact with DNA is essential for understanding gene regulation. While DNA-binding specificities for thousands of transcription factors (TFs) have been determined, the specific amino acid-base interactions comprising their structural interfaces are largely unknown. This lack of resolution hampers attempts to leverage these data in order to predict specificities for uncharacterized TFs or TFs mutated in disease. Here we introduce rCLAMPS (Recognition Code Learning via Automated Mapping of Protein-DNA Structural interfaces), a probabilistic approach that uses DNA-binding specificities for TFs from the same structural family to simultaneously infer both which nucleotide positions are contacted by particular amino acids within the TF as well as a recognition code that relates each base-contacting amino acid to nucleotide preferences at the DNA positions it contacts. We apply rCLAMPS to homeodomains, the second largest family of TFs in metazoans, and demonstrate that it learns a highly effective recognition code that can predict *de novo* DNA-binding specificities for TFs. Furthermore, we show that the inferred amino acid-nucleotide contacts reveal whether and how nucleotide preferences at individual binding site positions are altered by mutations within TFs. Our approach is an important step towards automatically uncovering the determinants of protein-DNA specificity from large compendia of DNA-binding specificities, and inferring the altered functionalities of TFs mutated in disease.

## 1 Introduction

Protein-DNA interactions are critical for the proper functioning of a wide range of biological processes within cells. Central among them is the regulation of gene expression, which is orchestrated by a complex network of sequence-specific interactions made by transcription factor (TF) proteins. While proper gene regulation ensures that the appropriate genes are expressed in each spatiotemporal context, mutations— within either TFs themselves or the particular genomic regions that they are intended to interact with—can alter gene expression programs and lead to disease phenotypes (review, Lee and Young (2013)). Due to the central importance of TF-DNA interactions in both healthy and disease states, there have been substantial efforts to characterize the determinants of specificity in TF-DNA interactions (review, Inukai *et al*. (2017)), and to catalogue DNA-binding specificities (Jolma *et al*. 2010; Berger and Bulyk 2009; Noyes *et al*. 2008a; Yang *et al*. 2017) and genomic occupancies (ENCODE *et al*. 2012; Moore *et al*. 2020) of TFs.

To date, DNA-binding specificities and/or context-specific genomic binding regions for thousands of TFs across human and other model organisms have been determined (Weirauch *et al*. 2014; Kulakovskiy *et al*. 2018; Mathelier *et al*. 2016b). These data have enabled large-scale construction of regulatory networks and have revealed the organization of regulatory circuits (Gerstein *et al*. 2012). Further, these data have been used to train machine learning models to uncover cooperative TF binding and other regulatory “grammars” (Avsec *et al*. 2021; Miraldi *et al*. 2021) and to predict the impact of mutations within noncoding regions of the genome (Zhou and Troyanskaya 2015; Martin *et al*. 2019). However, despite these extensive catalogues of DNA-binding specificities for wildtype TFs, as yet very few methods have been able to use these data in order to uncover the underlying determinants of protein-DNA specificity; this prevents both the prediction of *de novo* specificities for uncharacterized TFs (e.g., including those in non-model organisms or those that have proved difficult to study experimentally) as well as the prediction of altered DNA-binding specificities due to mutations or SNPs within TFs (e.g., as observed in several Mendelian disorders (Veraksa *et al*. 2000), cancers (Kobren *et al*. 2020), and across healthy populations (Barrera *et al*. 2016)). The main challenge is that for the vast majority of TFs for which DNA-binding specificities are known, the specific amino acid-base interactions comprising the underlying interaction interfaces are largely unknown, thus making it difficult to infer base preferences for specific amino acids or to determine which positions within their binding sites, if any, would be altered by mutations at specific amino acid positions.

Here, we introduce rCLAMPS (Recognition Code Learning via Automated Mapping of Protein-DNA Structural interfaces), a general probabilistic approach that uses DNA-binding specificities for TFs from the same structural family to simultaneously infer both which nucleotide positions are contacted by particular amino acids within the TF as well as a recognition code that relates base-contacting amino acids to nucleotide preferences at the DNA positions they contact. Our approach leverages the fact that protein-DNA interactions can be classified into structural families (Luscombe and Austin 2000) and that proteins from the same structural family have relatively well-conserved structural interfaces, where analogous positions within TFs tend to interact with the same set of DNA positions and these pairwise amino acid-base interactions define a contact map. rCLAMPS takes as input a collection of position weight matrices (PWMs) representing DNA-binding specificities for TFs from the same structural family, and uses a contact map representation of their protein-DNA structural interface. It performs Gibbs sampling to infer for each TF the mapping from its PWM columns to DNA positions within the contact map (i.e., determining which PWM columns across the TFs are analogous to each other) while simultaneously learning the parameters of a linear model that describes the base preferences of amino acids in specific positions of the TFs. Our approach is general and can be applied to PWMs for any family of DNA-binding proteins whose interaction interfaces with DNA are structurally conserved enough to be modeled by a pairwise amino acid-to-base position contact map.

We show the efficacy of our method by applying it to homeodomains, members of which play critical roles in development and cell fate processes (Burglin and Affolter 2016), and mutations within which are associated with a plethora of diseases (Chi 2006). We demonstrate that rCLAMPS accurately identifies analogous positions across a diverse set of homeodomain PWMs by mapping TF-PWM pairs to a canonical homeodomain contact map. Additionally, we show via extensive testing that rCLAMPS infers a recognition code that has excellent performance in predicting *de novo* DNA-binding specificities for homeodomain proteins. Finally, because rCLAMPS identifies which base positions within a PWM are contacted by specific amino acids within a TF and relies on an underlying linear model, we transfer existing specificity information from wildtype TFs to mutant TFs while making predictions only for the affected base positions, and demonstrate that this improves the accuracy of predicted specificities for mutant TFs beyond that of *de novo* predictions. Overall, we establish that our probabilistic framework yields a general, effective, and interpretable method that newly enables fully automated analyses of large compendia of known DNA-binding specificities in order to uncover protein-DNA interaction interfaces, infer recognition codes, and characterize mutant TFs.

### Further related work

Due to the fundamental importance of sequence-specific protein-DNA interactions in gene regulation, the determinants of TF-DNA interaction specificity have been studied extensively from both structural and statistical perspectives (Luscombe and Austin 2000; Luscombe 2001; Rohs *et al*. 2009; Persikov and Singh 2011). Machine learning methods to predict *de novo* specificities for TFs have been developed for several structural families, including Cys2His2 zinc fingers and homeodomains. Typically, these approaches to infer recognition codes explicitly incorporate prior knowledge of known DNA-contacting residues or curated protein-DNA interface contact maps (Noyes *et al*. 2008b; Benos *et al*. 2002; Kaplan *et al*. 2005; Persikov *et al*. 2009; Persikov and Singh 2014; Gupta *et al*. 2014; Persikov *et al*. 2015; Najafabadi *et al*. 2015). That is, the pairwise contacts between specific amino acids in the protein sequence and the DNA-binding sites are often known via specialized experimental protein-DNA interaction assays that position and orient the TF relative to a potential DNA binding site in a predetermined way (Noyes *et al*. 2008b,a; Chu *et al*. 2012; Persikov *et al*. 2014; Najafabadi *et al*. 2015); in contrast, our approach does not require *a priori* knowledge of these contacts and instead simultaneously infers them while learning a probabilistic recognition code. Alternatively, PWMs representing binding specificities for TFs from the same structural family have been aligned and subsequently machine learning models have been trained on them to predict DNA-binding specificities, an approach taken by the earliest method for homeodomains (Christensen *et al*. 2012); however, since the PWMs for TFs from the same structural family may be quite varied, these types of approaches require some manual intervention whereas rCLAMPS learns these alignments automatically. Protein-nucleic acid recognition codes have also been inferred from unaligned DNA sequences (Pelossof *et al*. 2015); while elegant, this approach relies on relating k-mers to quantitative measurements and these are only available for some experimental assays, whereas rCLAMPs uses PWMs and can be applied to binding specificites determined from any technical platform. Additionally, a model for predicting DNA-binding preferences for zinc finger proteins has been learned from unaligned DNA sequences (Kaplan *et al*. 2005); however, unlike rCLAMPS, this approach uses a manually curated contact map and assumes that nucleotide preferences are largely determined by single amino acid positions. Finally, several computational approaches have investigated large compendia of TFs and their corresponding DNA-binding specificities to learn rules for when the specificity of a characterized TF may be transferrable to an uncharacterized TF of the same family (Weirauch *et al*. 2014; Lambert *et al*. 2019); while powerful, these approaches cannot predict how a mutation within a TF will change its specificity. In contrast to previous approaches, we leverage structural information to enable automated, simultaneous inference of protein-DNA interaction interface mappings *and* an interpretable family-level DNA recognition code; consequently our approach enables us to determine when and how transfer of specificity information from wildtype to mutant TFs is appropriate at the resolution of individual binding site positions.

## 2 Methods

### 2.1 Overview of approach

Our framework rCLAMPS takes a corpus of DNA-binding specificities for a set of proteins from the same DNA-binding family; these DNA-binding specificities are represented as PWMs. It also utilizes protein-DNA co-complex structural data for that DNA-binding family in order to determine conserved pairwise contacts between positions in the proteins and positions within DNA that together comprise a structural interface or “canonical” contact map (Figure 1, top middle). For all pairs of TF proteins and their corresponding PWMs, the amino acids that correspond to each of the base-contacting positions within the contact map are known; however, the positions within the PWM that are contacted by these amino acids are not known (Figure 1, left). Our framework assumes that for each base position within the contact map, the base preferences at that position can be described by a linear model with additive contributions from amino acids occupying protein positions that contact that base position; each base position in the model is thus allowed to be involved in interactions with multiple amino acid positions. We use a linear model as its correspondence with the contact map makes it readily interpretable, especially for characterizing the impact of amino acid mutations on DNA-binding specificities. Initially, the parameters of the model are not known. However, if the parameters were known, then the likelihood of the data given each possible mapping of PWM columns to the nucleotide positions within the contact map can be computed. rCLAMPS uses a Markov chain Monte Carlo Gibbs sampling approach (Figure 1, bottom middle) to simultaneously learn the parameters of the model *and* determine which columns within the PWMs map to the same base position in the contact map (i.e., which columns across the PWMs are analogous to each other, Figure 1, right).

**Figure 1:**
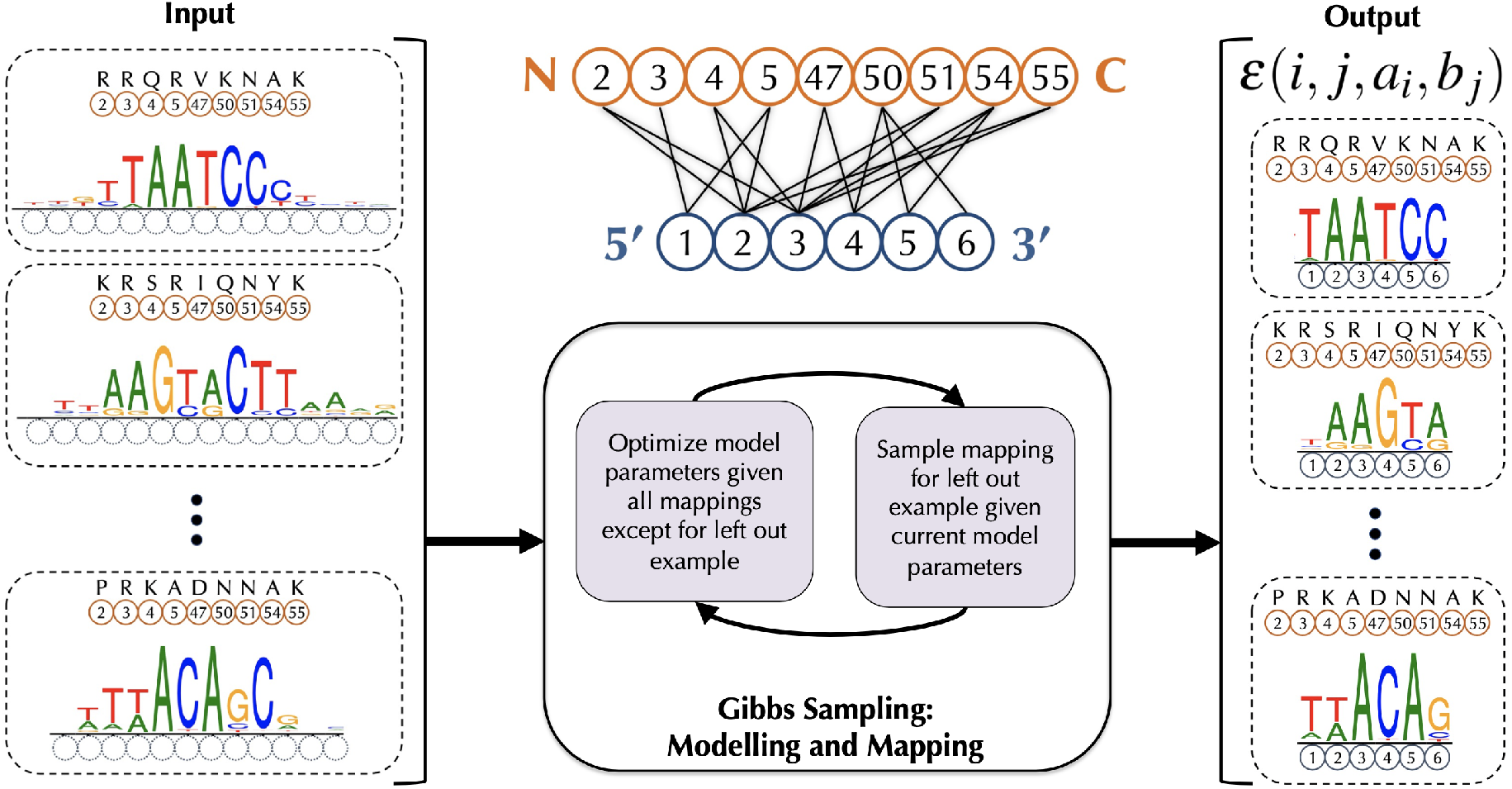
Schematic of our procedure rCLAMPS for jointly learning protein-DNA interaction interfaces and structure-aware recognition codes for TFs of the same structural family. **(Middle, top)** Our approach first analyzes protein-DNA co-complex structural data for a TF family to determine commonly observed pairwise contacts between positions in the protein (orange circles) and positions within DNA (blue circles) that together comprise a structural interface or “canonical” contact map. Here we show such a contact map for the homeodomain TF family, with protein positions corresponding to match states in Pfam homeodomain model PF00046. **(Left)** Given a set of TFs and their corresponding DNA-binding specificities as PWMs, the positions (and amino acids) within each TF that interact with DNA are known (orange circles and amino acids above), but initially the positions within the PWMs that are contacted by these amino acids are not known (dotted blue circles). **(Middle, bottom)** We use a Gibbs sampling approach to map the PWM positions to DNA positions within the contact map wherein base preferences at each nucleotide position are described in terms of additive amino acid-to-base contact energies. **(Right)** After Gibbs sampling is complete, we have a mapping of each TF-PWM pair to the TF family contact map, along with a linear recognition code for the TF family that consists of pairwise energy estimates for each amino-to-base pairing in each of the (*i, j*) amino acid-to-nucleotide position pairs in the contact map.

### 2.2 Deriving a contact map representation of the protein-DNA structural interface

For a DNA-binding domain of interest (i.e., a structural family of TFs), we obtain from BioLip (Yang *et al*. 2013) all co-complex structures of proteins from that family interacting with DNA. For each protein, we run HMMer with a Pfam model for that DNA-binding domain to map protein positions to HMM match states (Finn *et al*. 2014), thus finding analogous protein positions across the TFs. For each co-complex structure, for each of its amino acids that corresponds to an HMM match state, we then find DNA bases contacted by it, defining a contact as a pair of non-hydrogen atoms between an amino acid side chain and a DNA base at a distance of at most 3.6Å. We perform a co-complex structural alignment based on this set of contacts to ascertain which nucleotide positions across structures map to each other (see Wetzel and Singh (2020) for additional algorithmic and co-complex structure processing details). This structural alignment allows computation of an aggregate contact frequency matrix *D* where *D*[*i, j*] corresponds to the sequence-weighted (Henikoff and Henikoff 1994) fraction of times across the structures that the amino acid in match state *i* is in contact with the nucleotide in structurally-aligned binding site position *j*. Without loss of generality, we index the binding site positions 5’ to 3’ for the strand that is more frequently contacted by the proteins.

A contact map *C* (i.e., a set of (*i, j*) match state-to-binding site position contact pairs) for the TF family is then constructed based on *D* in the following way: We first identify a set of base-contacting match states based on a contact frequency threshold *t_d_*, 0 < *t_d_* < 1; match state *i* is included if in at least *t_d_* of sequence-weighted aligned co-complex structures the amino acid in match state *i* contacts any base. For each match state *i* that is base-contacting, we then add contact pair (*i, j*) to *C* if *D*[*i, j*] ≥ *t_e_*,0 < *t_e_* < 1. Note that the contact map represents a consensus “canonical” representation of which amino acid and nucleotide position pairs tend to be observed to be in contact across protein-DNA co-complex structures; a co-complex structure need not contain all contacts included in the map. For the homeodomain contact map derived here from 73 BioLip structures, we set both *t_d_* and *t_e_* equal to 0.05. This results in a contact map including eight contiguous base positions. We ultimately reduce this contact map to only the central six base positions of this contiguous region, which corresponds to the “core” homeodomain binding site illustrated in (Noyes *et al*. 2008b). These eliminated flanking positions of the homeodomain binding sites display on average lower information content in their DNA-binding specificities (see, e.g. Figure 2 of (Christensen *et al*. 2012), labeled as positions 1,2 and 9) and less consistency across repeated experiments for the same protein.

**Figure 2:**
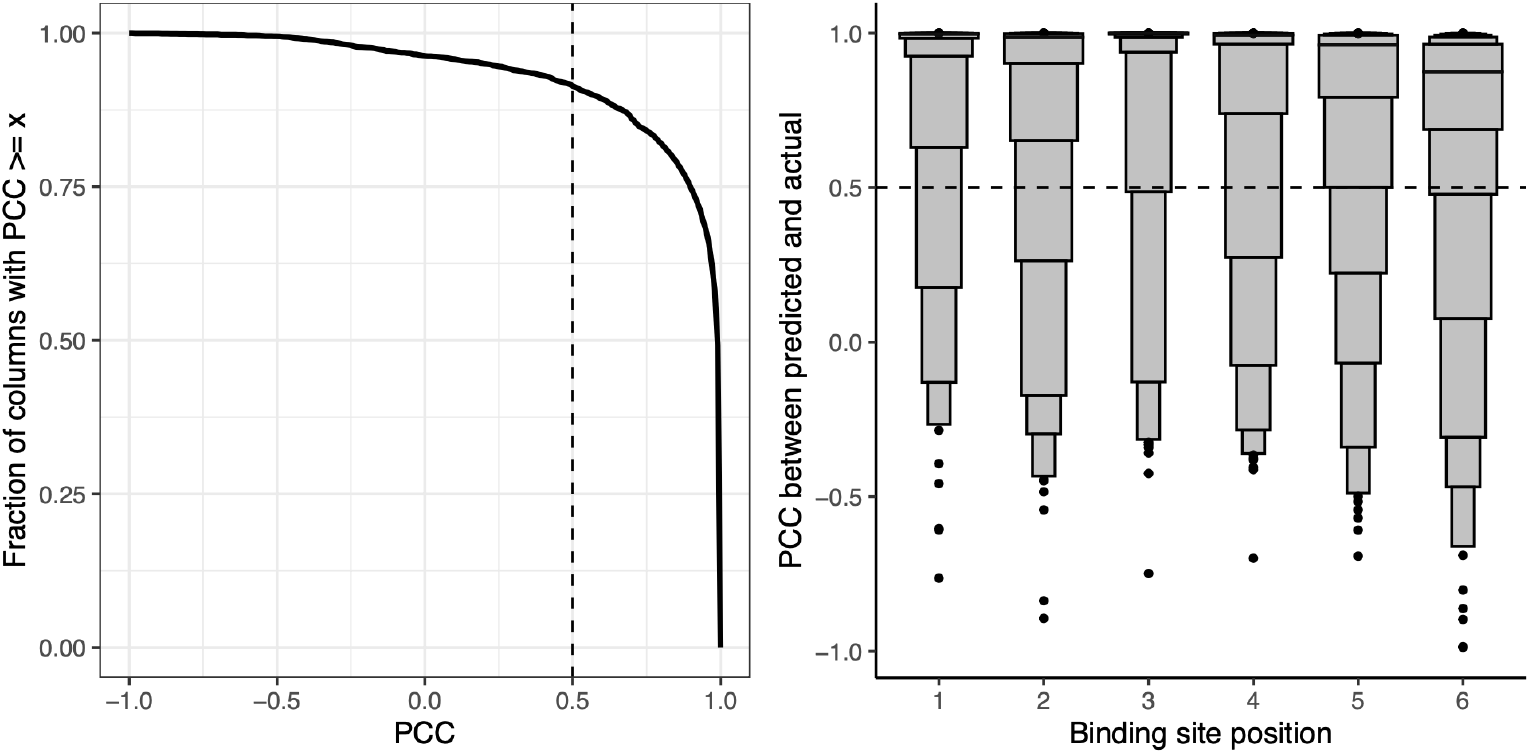
Probabilistic DNA recognition code for homeodomains derived from automatically inferred structural mappings has excellent *de novo* predictive performance. We compare agreement between predicted PWM columns and corresponding experimental PWM columns for 763 homeodomain proteins in a strict holdout validation setup. **(Left)** Considering all homeodomain binding site positions together, as different thresholds of PCC are considered (*x*-axis), the fraction of column pairs that have PCC greater than this threshold is plotted (*y*-axis). Our nominal threshold for agreement (PCC ≥ 0.5) is shown as a dashed vertical line. **(Right)** For each binding site position within the homeodomain contact map (*x*-axis), we display the PCC agreement scores (*y*-axis) for the paired columns at that binding site position, visualized as letter-value (or boxen) plots. In a letter-value plot, the widest box shows the value range spanned by half the data (from the 25th to 75th percentiles), while each successively narrower pair of boxes together show the value range spanned by half the remaining data. The PCCs at the 25th percentile for positions 1-6 are 0.98, 0.90, 1.00, 0.97, 0.79 and 0.69, respectively.

### 2.3 Model representation

For a particular protein-DNA interaction interface, we consider a model where the free energy of binding is additive over amino acid-binding site position pairs that are in the contact map *C*. That is, if *a_i_* is the amino acid in base-contacting position *i* and *b_j_* is the nucleotide in amino acid-contacting position *j*, and ***ε***(*i, j, a_i_, b_j_*) represents the energetic contribution arising from the contact between *b_j_* and *a_i_*, then the free energy of the entire protein-DNA interface is given by

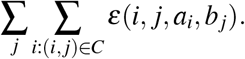

The energy contributions of each base *j* to the free energy of the entire protein-DNA interface is given by

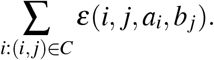

For a protein-DNA interaction interface, we assume that the identity of a base at any position *j* of the binding site is conditionally independent of the identity of bases at all other positions, given the amino acid sequence of the protein. For each amino acid position *i* and binding position *j* in the contact map, we introduce respectively random variables *A_i_* that can take on each of the 20 amino acids and random variables *B_j_* that can take on each of the four nucleotides. When considering a specific protein sequence (i.e., where *A_i_* = *a_i_* for all *i*), the Boltzmann distribution specifies the natural log of the probability that a particular base *b* is found in binding position *j* as

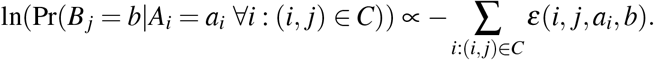

That is, if we know the amino acids occupying each position of a protein, we estimate the probability of a particular base at that *j*-th binding site position by considering just the sum of the pairwise binding energies of that base and the amino acids it interacts with according to our contact map *C*. Initially, these pairwise binding energies are not known, but in Section 2.5, we show how to estimate them from known protein-DNA structural interfaces.

### 2.4 Representations of DNA-binding specificities

For each TF, we assume that its DNA-binding specificity is represented as a PWM *F*, where entry *f_bj_* is the frequency with which nucleotide *b* is observed in column *j* in a set of aligned binding sites of that TF. Note that while binding sites for a single TF are aligned relative to one another, they are not yet mapped to our contact map *C*. That is, the analogous positions across a set of PWMs for a TF family are not typically known at the outset, and rCLAMPS will infer this. We transform each PWM to an estimated position count matrix (PCM) by simply multiplying each such frequency by 100 and rounding to the nearest integer. We note that our approach can also be applied to PCMs derived directly from aligned binding sites, but most commonly, specificity information for TFs are encapsulated in databases as PWMs. Furthermore, having the same number of total counts for each column of each PCM guarantees that each specificity in our training set contributes equally to the optimization procedure described below.

### 2.5 Estimating pairwise energy terms from mapped protein-DNA structural interfaces

Here we show that if we have a set of TFs and their PWMs along with a mapping of each TF’s PWM onto the structural interface (as in Figure 1, right), then we can estimate the pairwise energy terms of our linear model. We transform each PWM into a PCM as described above. Based on the Boltzmann distribution, for each base position *j*, we can relate amino acid outcomes to base probabilities via a log-linear model of the following form with one equation for each base outcome *b*:

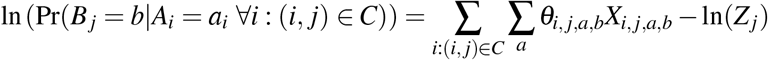

where *a* ranges over the 20 amino acids, *X_i,j,a,b_* is an indicator variable that is set to 1 if the amino acid in position *i* is *a, **θ**_i,j,a,b_* is the coefficient to be estimated that represents the contact energy contribution when nucleotide *b* in binding site position *j* is paired with amino acid *a* in protein position *i*, and *Z_j_* is a partition function. In particular, denoting parameters and indicator variables for the equation for base *b* in position *j* more compactly as 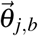 and 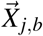, respectively,

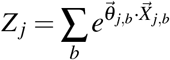

enforces ∑*_b_*Pr(*B_j_* = *b*|*A_i_* = *a_i_* ∀*i*: (*i, j*) ∈ *C*) = 1 for each combination of amino acid settings.

If we fix some an arbitrary base *b_o_* as a “reference” and in turn fix 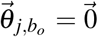, then each ***θ**_i,j,a,b_* can be interpreted as the expected contribution of a particular amino acid *a* at protein position *i* to a change in the energy of binding induced by swapping base *b* for base *b_o_* in base position *j*. Equivalently, since 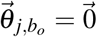, ***θ**_i,j,a,b_* is the expected change in log-odds of observing base *b* relative to base *b_o_*, given amino acid *a* at protein position *i*. Thus in practice we solve an equivalent multinomial logistic regression with the the `scikit-learn` Python package and regularize the model by adding pseudo-counts to the PCMs, corresponding to a flat Dirichlet prior. Ultimately ***ε***(*i,j,a_i_,b*) is set to the negative of its corresponding inferred model coefficient, since custom dictates that more favorable energetic states are more negative while more favorable log-odds are positive.

### 2.6 Mapping a PWM to the contact map using the energy terms

We next show that if we know the pairwise free energy binding terms, then we can compute the likelihood of each mapping of the TF’s PWM to the contact map. We are given a protein *p* with PWM *F_p_* = (*f_bj_*) and compute the corresponding PCM *K_p_* = (*k_bj_*). We want to infer *S_p_*, the index of the column of *F_p_* that maps to the first binding site position in the contact map *C*, as well as the orientation *O_p_* of the PWM with respect to the contact map. If *O_p_* = 1, then *F_p_* is mapped to *C*, and if *O_p_* = 0, the reverse complement of *F_p_* is mapped to *C*. For each possible setting of *O_p_* and *S_p_*, we compute the probability of observing the bases occupying the binding site positions in *C* in the PCM as follows. If *O_p_* = 1, and *w* is the number of nucleotide positions in our contact map *C*, then the sum of the natural logs of probabilities of observing the bases in the PCM is proportional to

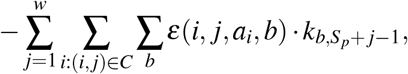

where *b* takes on each of the four nucleotides. On the other hand, if *O_p_* = 0, we let 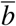 denote the base complementary to *b*, and then the sum of the natural logs of probabilities of observing the bases in the PCM is proportional to

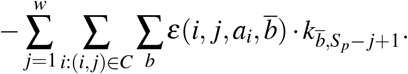

We refer to the set of mapping parameters for all proteins together as the registration *R* = {*R_p_*}, where *R_p_* = (*O_p_*, *S_p_*).

### 2.7 Gibbs sampling to estimate energy and mapping parameters

Initially, the pairwise contact energy terms ***ε*** are not known to us, and neither is the set of mapping parameters R. However, as we showed in Section 2.5, if the mapping is known, then all the ***ε*** terms can be estimated. On the other hand if the ***ε*** terms are known, then as we showed in Section 2.6, we can compute the probability of the observed data for each setting of the mapping parameters *R*.

Thus, we use Gibbs sampling for parameter inference. The Gibbs sampler initializes the mappings *R* randomly. During any given iteration of the sampling procedure, we hold out a protein *p* (and corresponding PCM *K_p_*), and estimate the ***ε*** terms as described in Section 2.5, using the the mapping *R_–p_* (i.e., withholding *R_p_* from *R*). We then sample a new *R_p_* proportional to the probability for each offset and orientation of *Kp* corresponding to a length *w* binding site based on these newly estimated *ε* terms, as described in Section 2.6. Sampling stops either when the joint probability of the data given the current settings of ***ε*** terms become bound in a small range over many iterations, or alternatively when a set maximum number of iterations is reached. In practice, we use a form of block sampling, where the mapping parameters for proteins with identical residues in their base-contacting positions are held out together and updated jointly (i.e., without recomputing the ***ε*** terms between members of the same blocks) in order to avoid being drawn into spurious local modes. Furthermore, we seed the sampling procedure with offset and orientation information for five protein-PWM pairs for which a co-complex structure already exists for the protein and the binding site in the structure aligns unambiguously to the corresponding PWM.

### 2.8 *De novo* prediction of PWMs

Once we have learned pairwise contact energy terms ***ε*** for a family of DNA-binding proteins as described in Section 2.5, then for a given protein *p* from that family, we first use Pfam to identify the amino acids within it that comprise the protein positions within the contact map. Next, for each binding site position *j* within the contact map *C*, we use

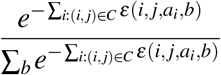

to calculate the probability with which each base *b* occurs in column *j* and set element *f_bj_* of a predicted PWM *F_p_* to this value.

### 2.9 Predicting PWMs for highly similar transcription factors

If we have two transcription factors that are nearly identical in the amino acids that comprise the structural interface, and we know the PWM for one of them and this PWM has been mapped to the structural interface, then the PWM for the other transcription factor can be predicted via a hybrid approach that transfers some PWM columns and predicts others *de novo*. In particular, given a TF *p* that is extremely sequence-similar to (i.e., is a mutant version of) another TF *p′*, for which we already know its PWM *F_p′_* and mapping *R_p′_*, we may transfer some information from *F_p′_* to infer a new PWM *F_p_*. Specifically, if the two proteins vary in a single DNA-contacting position *i*, then for each binding site position *j* that *does not contact* protein position *i* according to our contact map *C*, we simply set column *j* of *F_p_* equal to the column of *F_p′_* that maps to to binding site position *j* (i.e, according to *R_p′_*). In practice, since there may be many such near-mutant proteins (like *p′*) for each given binding site position *j* we use a weighted column transfer of the mapped column *j* across all such proteins, weighting each column by the corresponding near-mutant protein’s overall fraction of DNA-binding domain identity to *p* (considering only match positions and employing a minimum identity threshhold of 0.8). On the other hand, for each column *j* where no qualifying near-mutant protein exists in our dataset of mapped TF specificities, we simply predict column *j* of *F_p_ de novo* as described in Section 2.8.

### 2.10 Homeodomain PWM and protein dataset

PWMs for each of 623 wildtype homeodomains TFs were extracted from the Cis-BP database (build 2.00) (Weirauch *et al*. 2014), considering only specificities for which a single homeodomain was the only DNA-binding domain in the protein, and using the PWM from the most recent publication. Additionally, PWMs for 151 synthetic homeodomains from (Chu *et al*. 2012) and 30 mutant homeodomains from (Barrera *et al*. 2016) were added to our dataset. After removing PWMs corresponding to proteins missing DNA-contacting HMM match states, we had PWMs for a total of 763 distinct proteins. Prior to training our models, we rescaled PWMs to account for differences in information content across datasets (using a method described in (Christensen *et al*. 2012)). Proteins corresponding to each PWM were downloaded from Uniprot (The Uniprot Consortium 2021) (using the longest isoform for each) and mapped to HMM match states using HMMer v.3 with the PF00046 HMM (Finn *et al*. 2014).

### 2.11 Evaluating agreement between PWMs for the same protein

We compare two PWMs for the same protein based on the Pearson correlation coefficient (PCC) of their corresponding columns when mapped to the binding site positions of our contact map *C*. For predicted PWMs, this mapping is known. For experimental PWMs where this mapping is not known, we use the set of contiguous experimental PWM columns that best align to the predicted PWM using a previously described method (Persikov and Singh 2014). PCC is particularly suitable for many of our analyses due to its insensitivity to information content differences of PWMs across datasets.

## 3 Results

We demonstrate the effectiveness of our framework to simultaneously learn probabilistic recognition codes and the contacts comprising protein-DNA interaction interfaces from compendia of DNA-binding specificities by applying it to homeodomain TFs, which constitute the second largest family of TFs in metazoans.

### Accurate *de novo* prediction of PWMs via a structure-aware recognition code

We ran rCLAMPS with the homeodomain contact map and a set of 763 homeodomain proteins along with their DNA-binding specificities. Since models explored by Gibbs sampling can be sensitive to starting parameters, we ran Gibbs sampling 100 times and considered the mapping *R* of PWMs to our contact map *C* with the highest observed likelihood score.

We first demonstrate that the inferred mappings of the PWMs can be used to yield recognition codes that are highly effective in predicting the DNA-binding specificities of held out homeodomain proteins. That is, for each protein *z*, we reestimate the energies ***ε*** while withholding that protein and any other protein with identical DNA-contacting residues, and then predict the protein’s DNA-binding specificity *de novo* as described in Section 2.8. Because we are using the inferred recognition code to make a prediction, the mapping of each column in this predicted PWM to the contact map *C* is known. We compare this predicted PWM to the corresponding experimental PWM (also mapped to *C*, using the mapping *R_z_*). Remarkably, over all TFs z, more than 91% of the predicted columns are in agreement with the corresponding actual columns (i.e., have PCCs ≥ 0.5) (Figure 2, left).

Considering each binding site position separately, the median per-column PCCs when comparing predicted and experimentally measured columns are 1.0, 0.99, 1.0, 0.99, 0.96, and 0.88 for columns 1–6 respectively (Figure 2, right). We find that 95%, 91%, 97%, 91%, 88% and 86% of the predicted columns in positions 1–6, respectively, agree with the actual columns. As expected based on the determinants of homeodomain specificity, the strongest agreement is observed for base position 3, corresponding to a highly conserved asparagine to adenine contact. Columns also agree remarkably well for positions 4, 5, and 6, which are among the most variable positions of the core binding site for naturally occurring homeodomains, and for which our dataset contains synthetic homeodomains explicitly designed to vary specificity in these positions (Chu *et al*. 2012). Thus, even though the amino acid-to-base contacts comprising structural interfaces are automatically inferred, models trained assuming that these structural interfaces are correct have excellent predictive performance. Moreover, our structure-aware approach is expressive enough to describe a highly accurate recognition code for homeodomains, yet constrained enough to allow excellent generalization for predicting novel homeodomain specificities.

### Linear approach is competitive with state-of-the-art combinatorial models

We compare the predictive performance of rCLAMPS to that of existing state-of-the-art methods for predicting homeodomain DNA-binding specificities. In particular, we consider two random forest approaches, one that was trained using naturally occurring homeodomains (`rf_extant` (Christensen *et al*. 2012)), and another that demonstrated the utility of incorporating synthetic homeodomain training data (`rf_joint` (Chu *et al*. 2012)). For both of these approaches, the dataset of TF-PWM pairs required heuristics with semi-manual algorithmic modifications to uncover corresponding base positions across the PWMs; this was followed by feature selection before a random forest was trained for each base position. To provide a fair comparison of *de novo* predictive performance across methods on diverse proteins, we reserve half of the synthetic (at random) and all of the mutant homeodomain proteins as a test set and rerun rCLAMPS on the remaining 593 pairs that do not overlap these reserved proteins in terms of DNA-contacting residue combinations. Of the 141 proteins unseen by either rCLAMPS or `rf_extant`, our model predicts 90% of columns correctly, 5% more than `rf_extant`. Notably, rCLAMPS displays greater accuracy for five out the six core binding site positions for homeodomains, with the most marked improvements in positions 3 through 6 (data not shown). Our comparison to `rf_joint` is more limited, as the 77 proteins not in either method’s training set cover only 16 distinct DNA-binding residue combinations. Nonetheless, on this limited test set, both methods predict 94% of columns accurately. While `rf_joint` predicts one more column correctly in each of binding site positions 2 and 4, our model predicts two more columns correctly in position 5. Thus, despite the fact that these previous approaches used combinatorial models that required hyper-parameter tuning to avoid overfitting, as well as extensive preprocessing for alignment of PWMs and homeodomain position feature selection (Christensen *et al*. 2012), our automated probabilistic approach requiring only interpretable linear parameter fitting provides comparable *de novo* predictive power.

### Learning structural mappings allows effective transfer of specificity information from wildtype to mutants TFs

While *de novo* DNA-binding specificity prediction is necessary for predicting binding preferences of completely uncharacterized TFs, in the context of natural variation and disease, there is great interest in predicting the changes in DNA-binding specificities induced via single-amino acid alterations to wildtype TFs. Mutated TFs are seen in cancer (Kobren *et al*. 2020), and in inherited diseases (Hamosh *et al*. 2005; Chi 2006). If such an alteration occurs in a DNA-contacting residue, as a first approximation, *de novo* prediction is necessary only for the binding site positions contacted by that altered residue, while the specificity for the remaining binding site positions can be transferred directly from the wildtype. The primary obstacle preventing such a “hybrid” approach is that for the vast majority of TFs, the underlying amino acid-base contacts involved in the interaction are unknown. However, since rCLAMPS infers *both* a structural mapping of the underlying amino acid-base contacts *and* a per-binding site position *de novo* recognition code, we next employ such a hybrid inference approach.

To test our hybrid approach, we considered the 593 proteins given as input to rCLAMPS from the previous section as “wildtype” specificities and inferred specificities for each of 88 “mutant” proteins in the held out test set. A protein from the held out set is considered to be a mutant to a given wildtype protein if it is at least 80% identical to the wildtype throughout all positions in the match states of the DNA-binding domain and differs in exactly one DNA-contacting residue. Across these 88 mutants’ inferred specificities, our hybrid approach infers 353 of the columns via transfer from corresponding wildtype specificities, and requires *de novo* predictions for 175 columns. Comparing these inferred columns to their experimental counterparts, we find that 90% of the columns predicted *de novo* are accurate versus 93% for the transferred columns (Figure 3, left), suggesting a potential advantage to using the transfer approach when possible. Additionally, the transferred columns are more accurate than the identical column positions for the same proteins when predicted *de novo* by either rCLAMPS or by `rf_extant` (Figure 3, right). We note that the `rf_joint` method includes all but 19 of the mutant proteins in its training set (spanning only 8 distinct DNA-contacting residue combinations). On this extremely small and not very diverse set of proteins, both our transfer approach and `rf_joint` predict all columns accurately. Taken together, our results illustrate that the protein-DNA interface mappings inferred by our approach effectively enable transfer of wildtype homeodomain specificity information at the level of individual binding site positions and in turn allows inference of more accurate DNA-binding specificities for mutant homeodomains than *de novo* prediction alone.

**Figure 3:**
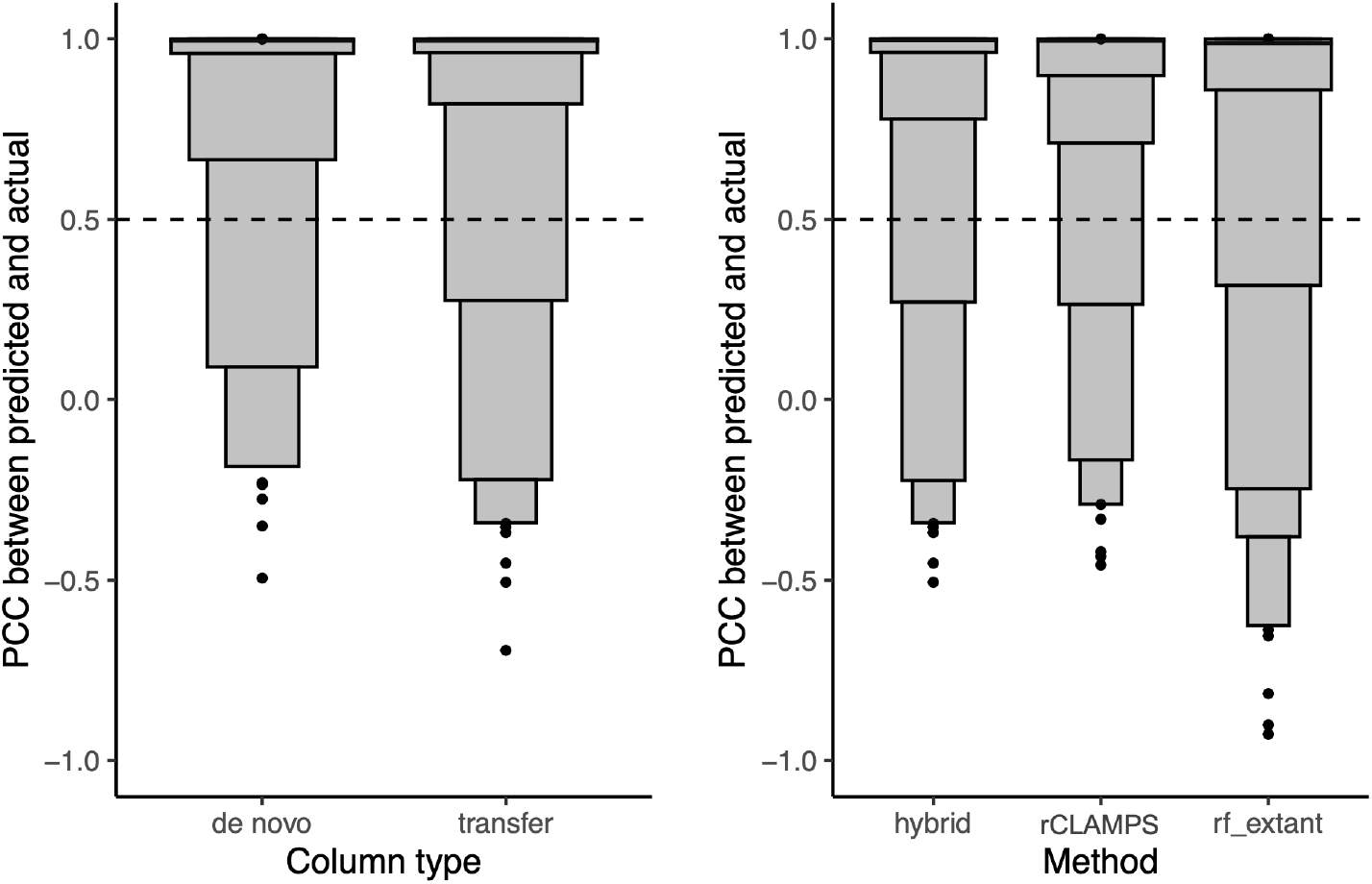
Column transfer via learned structural mappings is highly effective for predicting DNA-binding specificities for mutant TFs. **(Left)** For the hybrid approach, the PWM columns are binned according to whether they are predicted *de novo* or via transfer, and the PCCs of the predicted vs. actual columns are shown in letter-value plots. While 90% of *de novo* predictions are in agreement with their experimentally determined counterparts, an even higher 93% of predictions via transfer are in agreement. **(Right)** For each of the homeodomains that were also not part of the `rf_extant` training set, and considering only columns for which transfer was used by our hybrid approach, we compute the PCC between the actual specificity for a column and that predicted by our hybrid approach (hybrid), our *de novo* linear approach (rCLAMPS), and the `rf_extant` model (rf_extant). We find that 93% of transferred predictions are in agreement with their experimentally determined counterparts, as compared to 91% and 85% of *de novo* predictions for rCLAMPS and `rf_extant`, respectively.

### Accurate and automated mapping of TF-PWM pairs to a structural interface

Because homeodomain proteins and their DNA-binding specificities are highly conserved across organisms (Nitta *et al*. 2015), we next use an across-species approach to externally validate our mapping of TF-PWM pairs using previously determined known protein-DNA interfaces. Specifically, in the first large-scale assay of homeodomains in fruit fly, DNA-binding specificities were characterized in a specialized experimental system that specifically allowed for a global alignment (with known orientation) of all the DNA sequences selected by all the DNA-binding domains assayed (Noyes *et al*. 2008b). By manually aligning a single PWM from this set relative to the start of our contact matrix and shifting registrations of all others identically, we obtain an experimentally inferred mapping of each of these fly PWMs. Of 593 homeodomains for which rCLAMPS inferred structural mappings, 235 of them map to fly proteins characterized in this previous assay as they are identical in their base-contacting positions. Thus, we compare correspondences between the experimentally determined fly mappings and the mappings of their base-contacting-residue-identical counterparts in our dataset.

Strikingly, 97% (228/235) of our inferred mappings are identical to their experimentally determined counterparts (Figure 4, left, green). Further, after separating the homeodomain proteins according to “specificity groups” as determined by the authors of the experimental approach via hierarchical clustering of their DNA-binding motifs (Noyes *et al*. 2008b), we find that the mappings are either completely or nearly completely correct for 10 out of 11 of these diverse specificity groups (Figure 4, right, green), with poor agreement occurring for only the relatively small *Iroquois* group.

**Figure 4:**
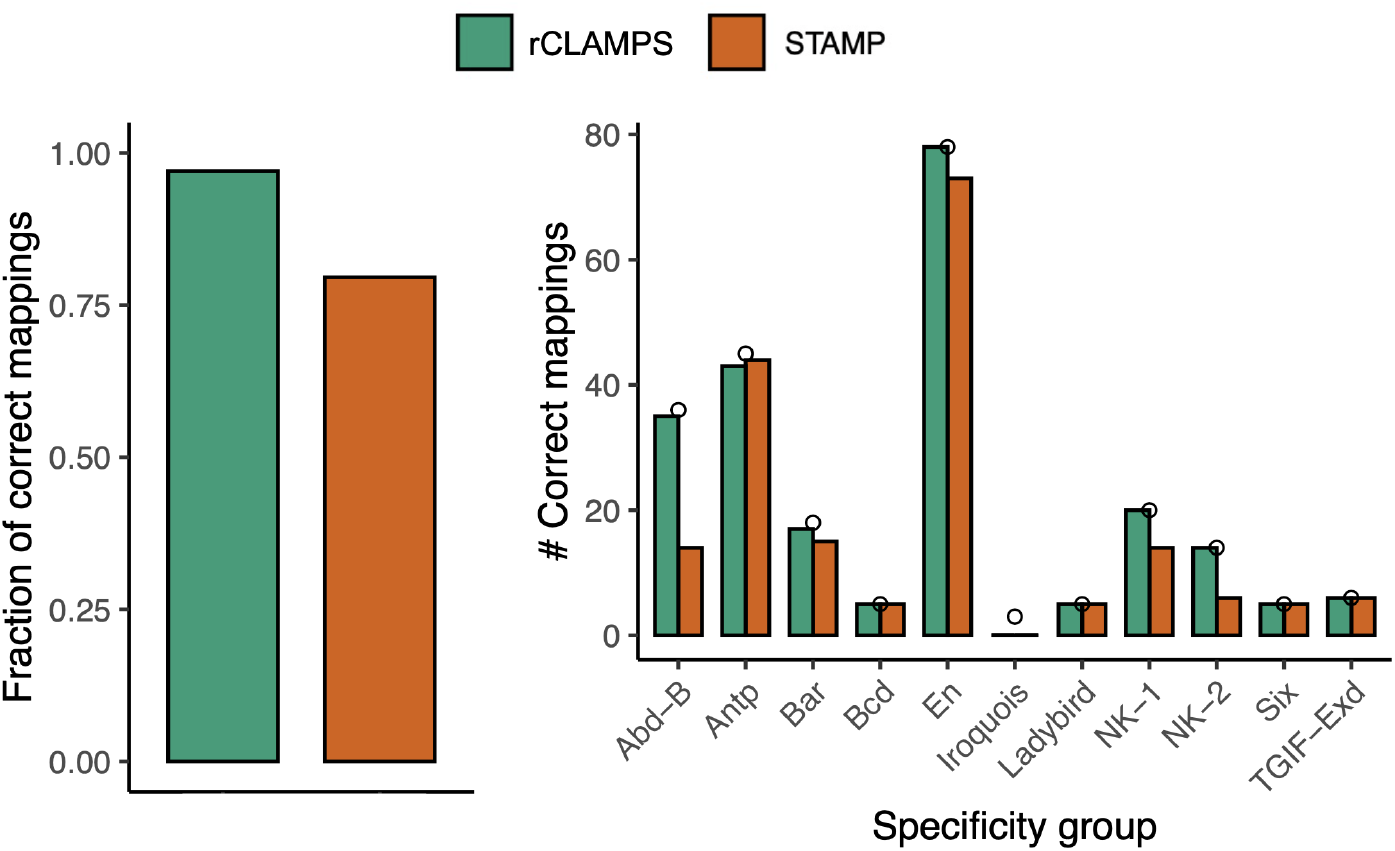
Inferred mappings to contact map are highly accurate. We compare mappings inferred by rCLAMPS (green) and computed based on direct PWM multiple alignment performed by STAMP (red) to those that are known experimentally for TFs that have identical amino acids within their base-contacting positions. **(Left)** The fraction of predicted mappings that are identical to the experimental mappings. **(Right)** The number of identical mappings, when homeodomains are classified with respect to ‘specificity groups’ as determined in (Noyes *et al*. 2008b). Small circles represent the total number of homeodomains considered in each of these specificity groups.

To illustrate the advantage of our approach that jointly considers the proteins and the binding sites when aligning specificities for a family of TFs, we compare our results to a tool that considers only binding site information. In particular, we consider STAMP (Mahony *et al*. 2007), which is a multiple PWM alignment tool that uses a guide tree based on pairwise PWM alignments (i.e., without considering the protein sequences associated with the PWMs). We run STAMP on the same set of PWMs and use default parameters. While STAMP does not map its aligned set of PWMs to the homeodomain contact map, we consider the mapping of the alignment which gives the best possible results for STAMP. This results in a correct mapping for 80% of the TFs (Figure 4, left, red). STAMP does well on the most abundant groups (e.g., *Antp* and *En*); this is as expected since these PWMs are similar to each other and align well to each other. In contrast, STAMP struggles on groups that diverge more substantially from the canonical homeodomain motif pattern (e.g. the *NK-2* group) or groups that tend to have reverse palindromic PWMs with more subtle specificity differences between the two sides of the palindrome (e.g. the *Abd-B* group). This is also as expected, since STAMP’s intended use is to align similar PWMs, not a diverse set of PWMs as can be seen for different TFs from the same structural family. Thus, for TF families with diverse or complex DNA-binding specificity determinants, our framework can effectively harness the shared information between the proteins and their corresponding PWMs to produce a multiple PWM alignment that is in outstanding agreement with a ground truth experimental alignment.

## 4 Discussion

We describe a novel probabilisitic framework that enables fully automated analyses of large compendia of TF DNA-binding specificities to jointly discover mappings of underlying sets of amino acid-to-base contacts and structure-aware TF family-wide recognition codes. In principle, our approach can be applied to PWMs for any family of DNA-binding proteins whose interaction interfaces with DNA are structurally conserved enough to be modeled by a pairwise amino acid-to-base position contact map. Using the homeodomain family as a test case, we show that the physically interpretable recognition code learned is both expressive enough and generalizable enough to allow state-of-the-art *de novo* prediction of homeodomain DNA-binding specificities. Furthermore, we demonstrate that having extremely accurate mappings of TF-PWM interfaces allows single base position resolution transfer of specificity information from wildtype to mutant proteins, in turn enabling inference of even more accurate DNA-binding specificities.

Our linear model and set of mapped TF-DNA interfaces can serve as a jumping off point for training models that account for higher order interactions, and without the need to rely on experimentally curated contact maps (Noyes *et al*. 2008b; Benos *et al*. 2002; Kaplan *et al*. 2005; Persikov *et al*. 2009; Persikov and Singh 2014; Persikov *et al*. 2015; Najafabadi *et al*. 2015), specialized experimental setups that place the protein in a fixed orientation with the binding site (Noyes *et al*. 2008b,a; Chu *et al*. 2012; Persikov *et al*. 2014; Najafabadi *et al*. 2015), or complicated and partly curated multiple motif alignment strategies (Christensen *et al*. 2012; Chu *et al*. 2012). Indeed, in a preliminary exploration where we trained machine learning methods that allow nonlinear effects between amino acids on base preferences (including gradient boosting (Friedman 2000) and random forests (Breiman 2001)) on our set of mapped TF-DNA interfaces, we found that they provide subtle improvements in predictive performance for some base positions. However, we do not report those results here as the pairwise energetic approximation provided by rCLAMPS allows for statistically interpretable coefficients while still providing state-of-the-art predictive performance. Optimizing these non-linear, more expressive models is likely to be a fruitful avenue for further research. Moreover, DNA shape has been shown to be an important determinant for both intrinsic and context-specific DNA-recognition by homeodomain and other TF families (Gordân *et al*. 2013; Dror *et al*. 2014; Zhou *et al*. 2015; Mathelier *et al*. 2016a; Yang *et al*. 2017; Kribelbauer *et al*. 2020); thus, extending our model to include DNA shape information based on binding sites’ flanking nucleotide contexts may lead to more accurate predictions of the effects of TF mutations on DNA-binding activity.

Overall, we expect that our general approach can be applied to the thousands of extant DNA-binding specificities across a range of organisms and to a diverse set of DNA-binding families; this would drastically improve our understanding of the determinants of DNA-binding specificity and in turn help efforts to predict the potential downstream regulatory impacts of mutations within TFs.

## 5 Data Access

Open-source software implementing methods and analyses described in this work are accessible at: https://github.com/jlwetzel-slab/rCLAMPS.

## 7 Acknowledgements

This research has been supported in part by NSF[ABI-1458457] and NIH[R01-GM076275].

